# Berberine Bridge Enzyme-Like Oligosaccharide Oxidases Act As Enzymatic Transducers Between Microbial Glycoside Hydrolases and Plant Peroxidases

**DOI:** 10.1101/2022.04.15.488465

**Authors:** Anna Scortica, Moira Giovannoni, Valentina Scafati, Francesco Angelucci, Felice Cervone, Giulia De Lorenzo, Manuel Benedetti, Benedetta Mattei

## Abstract

OG-oxidases (OGOXs) and CD-oxidase (CELLOX) are plant berberine bridge enzyme-like oligosaccharide oxidases that oxidize oligogalacturonides (OGs) and cellodextrins (CDs), cell wall fragments with nature of damage-associated molecular patterns (DAMPs). The oxidation of OGs and CDs attenuates their elicitor activity by concomitantly releasing H_2_O_2_. Here, we demonstrate that the H_2_O_2_ generated downstream of the combined action between a fungal polygalacturonase and OGOX1 or an endoglucanase and CELLOX can be directed by plant peroxidases (PODs) either towards a reaction possibly involved in plant defence such as the oxidation of monolignol or a reaction possibly involved in a developmental event such as the oxidation of auxin (IAA), pointing to OGOX1 and CELLOX as enzymatic transducers between microbial glycoside hydrolases and plant PODs.

## MAIN MANUSCRIPT BODY

Plants are constantly menaced by a wide array of pathogens. Against them, plants evolved a robust barrier composed of polysaccharides and phenolic compounds, i.e., the cell wall, and a sophisticated defence system that can be promptly activated at the occurrence. In order to colonize the plant tissue, pathogens need firstly to dismantle the cell wall, whose degradation is achieved through the secretion of cell wall degrading enzymes that include glycoside hydrolases (GHs), esterases and oxidoreductases (Benedetti et al., 2019; Giovannoni et al., 2020). The enzymatic hydrolysis of cell wall polysaccharides may result in the transient accumulation of cell wall fragments such as oligogalacturonides (OGs), cellodextrins (CDs) and other cell wall oligosaccharides that are quickly perceived by plants as danger signal, i.e., as damage-associated molecular patterns (DAMPs) (Pontiggia et al., 2020).

How plants modulate the amplitude of defences in response to the extent of cell wall hydrolysis is not known. Cell wall degradation occurs not only upon a microbial attack but is also necessary for remodelling during development. Therefore, cell wall fragments can also be generated by endogenous enzymes during the relaxation of the cell wall structures, pointing to the necessity of a system capable of discriminating an exogenous infection from an endogenous developmental stimulus. Thus, a system capable of measuring the entity of a cell wall damage must exist.

Some berberine bridge enzyme-like (BBE-l) proteins from *Arabidopsis thaliana* have been recently identified as specific OG-oxidases (OGOXs) and CD-oxidases (CELLOXs). OGOXs include four isoforms (OGOX1-4) encoded by paralogous genes that are capable of oxidizing galacturonic acid oligomers of different size (OGs), whereas CELLOX oxidizes CDs (Benedetti et al., 2018; Locci et al., 2019). Structural data of two Arabidopsis BBE-l Monolignol-oxidases (Daniel et al., 2015; Daniel et al., 2016) as well as 3D structural modeling and amino acid alignment of the four OGOXs, CELLOX and other plant BBE-l carbohydrate oxidases allowed to identify features important for oxidase activity including the residue V155/157 of OGOX1/CELLOX (Benedetti et al., 2018; Locci et al., 2019) as the gatekeeper residue of the oxygen binding pocket [P(T/S)VGVGG] (Leferink et al., 2009; Zafred et al., 2015). Indeed, OGOXs and CELLOX inactivate the elicitor nature of OGs and CDs by concomitantly releasing H_2_O_2_, a molecule with multiple functions in the cell wall strengthening and signalling (Smirnoff and Arnaud, 2019). The oxidized oligosaccharides are characterized by an increased recalcitrance to enzymatic hydrolysis (Benedetti et al., 2018), but nothing is known about their involvement in other physiological processes. Recently, the combined use of Arabidopsis OGOX1 and a peroxidase (POD) allowed the measurement of the OGOX1 activity suggesting that possible physiological processes could be driven by the OGOX-generated H_2_O_2_ in the presence of POD (Scortica et al., 2021). In the present study, the capability of generating H_2_O_2_ by combinations of OGOX1 with a microbial polygalacturonase and CELLOX with a microbial endoglucanase was tested. The generated H_2_O_2_ can be utilized as a substrate by POD for oxidative reactions possibly involved in defence and development (Fig. 1). Indeed, glycoside hydrolases (GHs), OGOX1, CELLOX and PODs perform their enzymatic function in the same cell compartment, i.e., the apoplast, and it is plausible to consider their activities as related in cell wall metabolism.

**Fig. 1.**
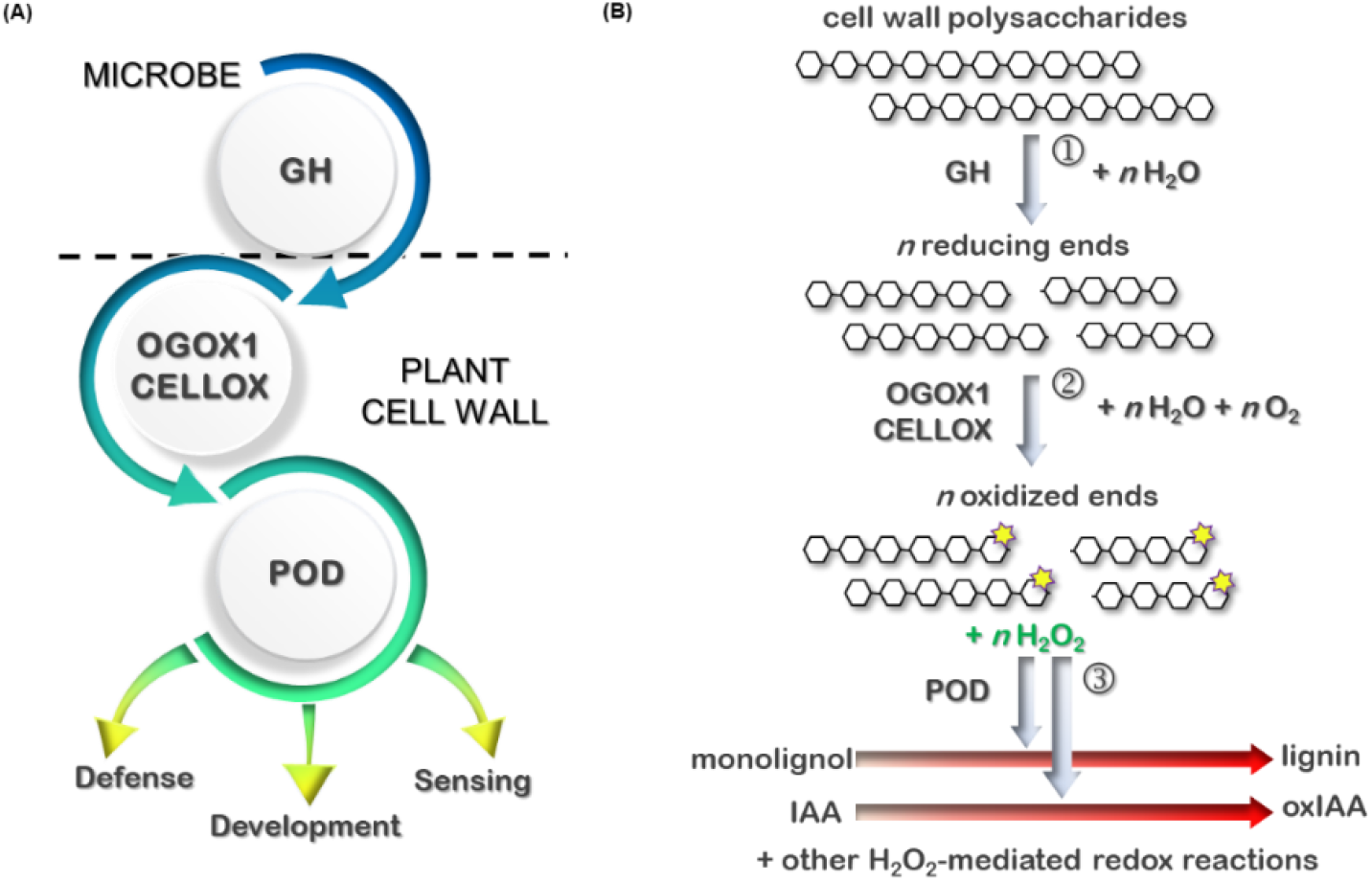
OGOX1 and CELLOX as transducers between microbial GHs and plant PODs. (A) Schematic representation showing the transducing role of OGOX1 and CELLOX between microbial GHs and plant PODs and their potential involvement in different plant processes. (B) Working model of a OGOX1/CELLOX-POD machinery: in step 1, microbial GHs hydrolyse the cell wall polysaccharides by generating reducing end-free oligomers. In step 2, specific BBE-l oligosaccharide oxidases (OGOX1 and CELLOX) oxidize such reducing ends by concomitantly releasing H_2_O_2_. In step 3, H_2_O_2_ is used by plant PODs to oxidize monolignols or IAA. [BBE-l: Berberine bridge enzyme-like, CELLOX: CD-oxidase from *A. thaliana*, GH: Glycoside hydrolase, OGOX1: OG-oxidase 1 from *A. thaliana*, POD: Peroxidase, IAA: indole-3-acetic acid, oxIAA: oxidized indole-3-acetic acid].

To evaluate whether, during a plant-microbe interaction, the combined activity of a plant-derived BBE-l oxidase and a microbial GH generates H_2_O_2_ that can be sequentially utilized by PODs to start biologically relevant reactions involved in defence and growth and therefore in the defence/growth trade-offs (Fig. 1), we used OGOX1 (Benedetti et al., 2018) and CELLOX (Locci et al., 2019) in combination with a recombinant endopolygalacturonase from *Fusarium phyllophilum* (FpPG) and a commercial endoglucanase from *Aspergillus niger* (AnEG), respectively. The commercial horseradish peroxidase VI-A type (HRP) that catalyzes the oxidative polymerization of guaiacol, here used as coniferyl alcohol analogue, and an anionic peroxidase preparation from ripe tomato fruit (APOD) that utilizes H_2_O_2_ to oxidize IAA (Kokkinakis and Brooks, 1979), a typical growth hormone, were used as representative plant PODs. FpPG, OGOX1 and CELLOX were expressed in *P. pastoris* and purified to homogeneity. The expression of OGOX1, was achieved as reported in (Scortica et al., 2021), whereas the expression of CELLOX, due to the low yield and high protein instability, required a different expression strategy that consisted in the addition of a Flag-6xHis-SUMOstar tag upstream of the sequence encoding CELLOX (Fig. S1A). The sequence encoding the sumoylated form of CELLOX (Data S1), here referred to as FHS-CELLOX, was cloned under the control of the methanol-inducible promoter AOX and expressed in *P. pastoris*. Immuno-decoration analysis performed on the culture filtrates from four different transformants showed that FHS-CELLOX is expressed in a heavily glycosylated form (Fig. S1B) and, upon de-glycosylation with PNGase F, appears as a unique polypeptide chain of 74 kDa (Fig. S1C). OGOX1 was purified from the culture filtrate of *P. pastoris* by two hydrophobic interaction chromatography steps performed at two different pH values (5.0 and 7.0) (Scortica et al., 2021), whereas FHS-CELLOX was purified in a single step by IMAC chromatography. The AnEG used in our experiments was a highly pure preparation from a commercial source whereas FpPG was constitutively expressed in *P. pastoris* and purified using a three-step purification procedure as reported in (Benedetti et al., 2011). The protein yields were about 5 mg.L^-1^, 0.5 mg.L^-1^ and 15 mg.L^-1^ for OGOX1, FHS-CELLOX and FpPG, respectively. Before proceeding with the enzymatic assays, the purity grade of the different protein preparations was assessed by SDS-PAGE/Coomassie blue staining analysis (Fig. S2). To evaluate the H_2_O_2_-conversion efficiency of OGOX1 and FHS-CELLOX, the amount of H_2_O_2_ released from the enzymatic oxidation of penta-galacturonic oligosaccharide and cello-triose, here used as model substrate of OGOX1 and FHS-CELLOX, respectively, was measured over reaction time. Our analysis clearly indicated that both OGOX1 and FHS-CELLOX are efficient reducing sugar-to- H_2_O_2_ converters, with H_2_O_2_ conversion efficiencies ranging from 85 to 95% (Fig. S3). Polygalacturonic acid and carboxy-methyl cellulose, respectively the substrates of FpPG and AnEG, were added to the two enzyme combinations FpPG-OGOX1-HRP and AnEG-(FHS-)CELLOX-HRP. In both reaction mixtures, HRP utilized the generated H_2_O_2_. As shown in Fig. 2, the degrading activity of FpPG and AnEG was quantitatively converted to tetra-guaiacol polymerization in a time-dependent manner, allowing to monitor the activity of both GHs over the entire reaction time. The absence of HRP or the BBE-l oligosaccharide oxidase or the addition of CAT in the reaction mixture prevented the tetra-guaiacol polymerization (Fig. 2). Taken together, these results indicated the central role of H_2_O_2_ as the molecule linking the activity of microbial GHs and plant PODs.

**Fig. 2.**
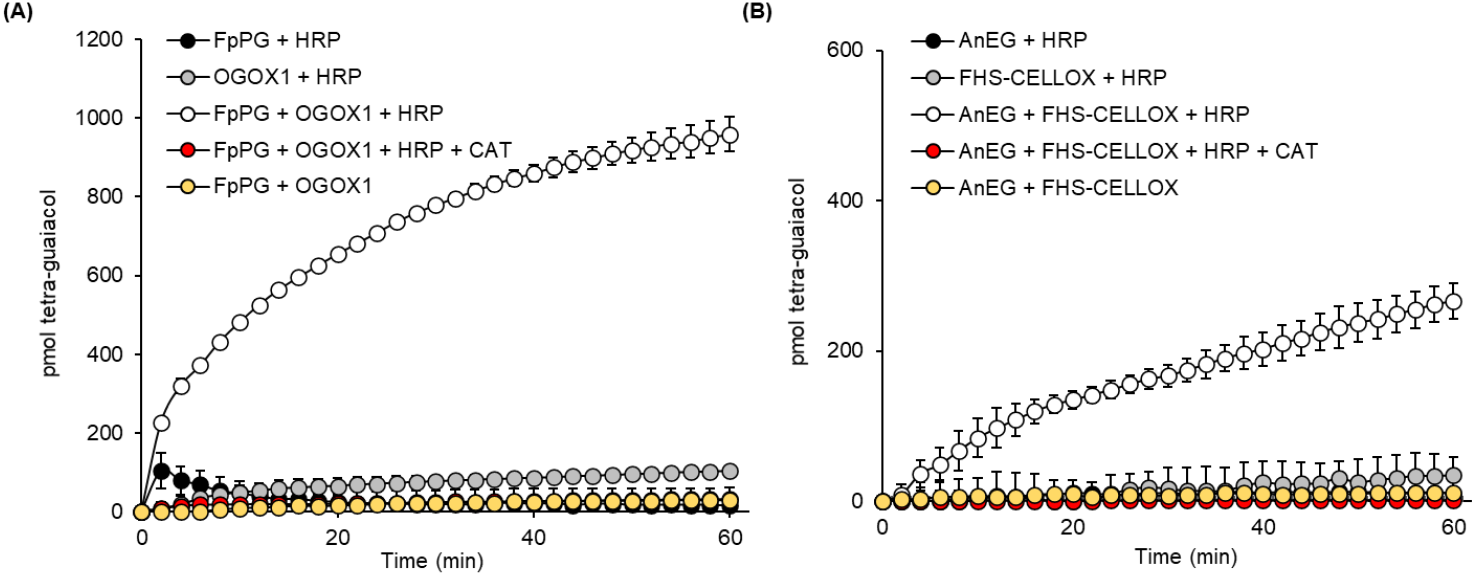
A OGOX1/CELLOX-POD machinery quantitatively converts a polysaccharide hydrolysis to tetra-guaiacol polymerization. (A, B) Tetra-guaiacol polymerization over time using a OGOX1/CELLOX-POD machinery formed by (A) FpPG, OGOX1 and HRP against polygalacturonic acid and (B) AnEG, FHS-CELLOX and HRP against carboxy-methyl cellulose. For each enzymatic machinery, different combinations of enzymes were used. As control, CAT was added to eliminate the H_2_O_2_ generated by each GH-OGOX1/CELLOX pair. Values are mean ± s.d. (n= 2). The experiments (A, B) were repeated twice with similar results. [AnEG: endoglucanase from *A. niger*, CAT: catalase from bovine liver, FHS-CELLOX: Flag-His-SUMOstar-tagged CD-oxidase from *A. thaliana*, FpPG: endopolygalacturonase from *F. phyllophilum*, OGOX1: His-tagged OG-oxidase 1 from *A. thaliana*, HRP: horseradish peroxidase VI-A type].

The ripe tomato fruit was used as source of APOD (Kokkinakis and Brooks, 1979). Before proceeding with the assays in combination with the OGOX1/(FHS-)CELLOX pairs, APOD activity was quantified using ABTS and H_2_O_2_ (Fig. S4). The same substrates, i.e., polygalacturonic acid and carboxy-methyl cellulose, were added to two enzyme combinations FpPG-OGOX1-APOD and AnEG-(FHS-)CELLOX-APOD, respectively. In both reaction mixtures, APOD utilized the generated H_2_O_2_. As shown in Fig. 3A-B, the degrading activity of FpPG and AnEG was quantitatively converted to oxidized IAA in a time-dependent manner, and activities of both GHs could be monitored by following the amount of residual (non-oxidized) IAA over reaction time. Also in this case, the lack of APOD or a BBE-l oligosaccharide oxidase prevented the IAA oxidation (Fig. 3A-B). These results taken together clearly demonstrate that the H_2_O_2_ generated downstream of the GH/BBE-l oligosaccharide oxidase pair is successfully used by plant PODs as oxidant in two different processes, i.e., tetra-guaiacol polymerization and IAA oxidation.

**Fig. 3.**
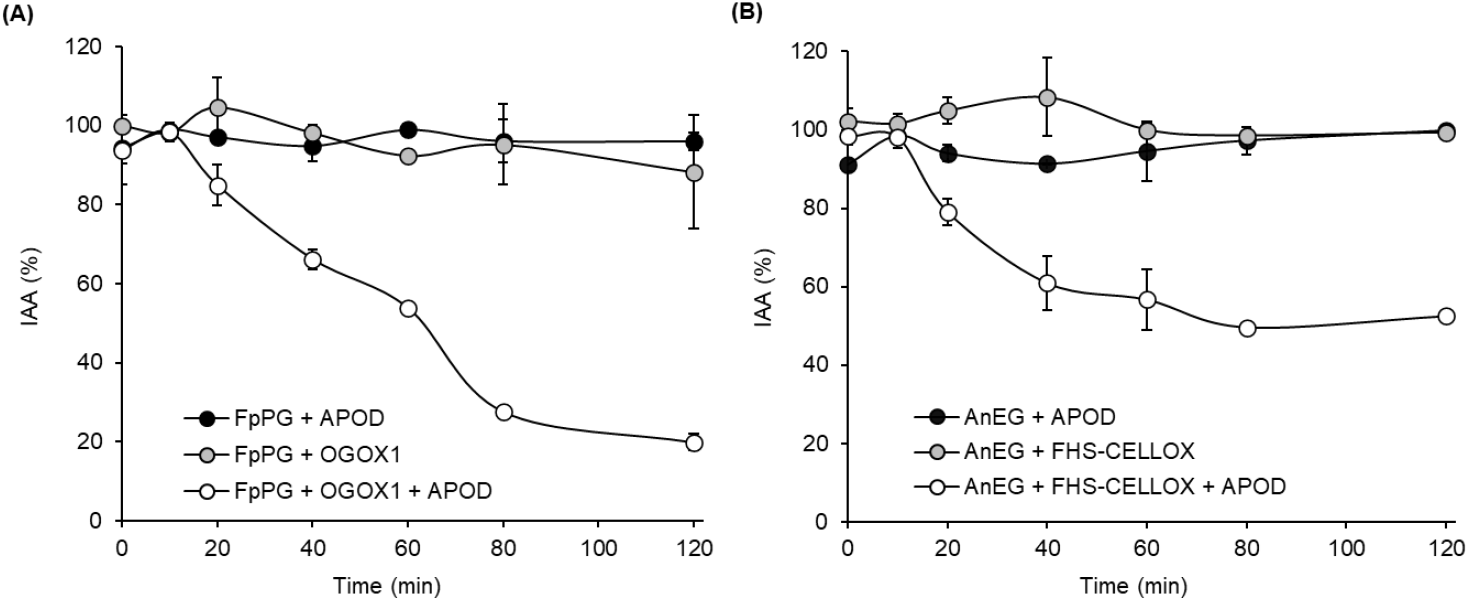
A OGOX1/CELLOX-POD machinery quantitatively converts a polysaccharide hydrolysis to IAA oxidation. (A, B) IAA oxidation over time using a OGOX1/CELLOX-POD machinery formed by (A) FpPG, OGOX1 and APOD against polygalacturonic acid and (B) AnEG, FHS-CELLOX and APOD against carboxy-methyl cellulose. For each enzymatic machinery, different combinations of enzymes were used. Values are mean ± s.d. (n= 2). The experiments (A, B) were repeated twice with similar results. [AnEG: endoglucanase from *A. niger*, APOD: anionic peroxidase preparation from ripe tomato fruit, FHS-CELLOX: Flag-His-SUMOstar-tagged CD-oxidase from *A. thaliana*, FpPG: endopolygalacturonase from *F. phyllophilum*, OGOX1: His-tagged OG-oxidase 1 from *A. thaliana*].

To date, OGOX1-4 and CELLOX are the only plant BBE-l proteins with proven oxidizing activities towards cell wall oligosaccharide fragments with elicitor nature, i.e., OGs and CDs. However, due to the large number of members constituting the different plant BBE-l families (Daniel et al., 2017; Pontiggia et al., 2020), it is plausible that other BBE-l enzymes still orphan of their substrate may act as specific oxidases of other cell wall-derived oligosaccharides. During the reaction catalysed by OGOXs and CELLOX, OGs and CDs are inactivated and H_2_O_2_ is formed. Unlike with other extracellular H_2_O_2_-producing enzymes such as the membrane bound NADPH oxidase (Kadota et al., 2015), H_2_O_2_ produced by OGOXs and CELLOX is produced only locally from the reducing end of OGs and CDs enzymatically liberated, either by an endogenous enzyme or, as in the case of a pathogenic attack, by microbial GHs at the site of infection where one molecule of H_2_O_2_ is generated from one free reducing end. During the degradation of the plant cell wall, the resulting OGs and CDs and possibly other cell wall fragments can be converted by OGOX and CELLOX and possibly other BBE-l oligosaccharide oxidases to H_2_O_2_ that, in turn, may be used by extracellular PODs to promptly reinforce the cell wall in a proportional opposite direction to the occurring degradation, i.e., more degradation is performed by microbes, more lignification occurs (Fig. 4). During the pathogen attack, the same enzymatic interplay may also cause inhibition of plant growth through an oxidation of the extracellular IAA (Fig. 4). The APOD-mediated oxidation of IAA could play a role in the growth-defence trade-off when plants are required to redirect their metabolic energy from primary to secondary metabolism during pathogen infection (Pontiggia et al., 2020). Thus, the type of molecule that will be oxidized by H_2_O_2_ will depend on the substrate specificity of the available plant POD. Considering that 73 different class III plant PODs exist in *A. thaliana* (Almagro et al., 2009) and that most of them are localized in the extracellular space, the oxidizing activity of H_2_O_2_ can be sorted towards several metabolic pathways. Indeed, Arabidopsis BBE-l oligosaccharide oxidases (OGOX1 and CELLOX) and several class III PODs are positively co-expressed during fungal infection, corroborating their involvement in a potential enzymatic interplay in plant defence (Fig. S5, Table S1). Our experiments clearly demonstrate that apparently unrelated enzymes such as glycoside hydrolases, the flavoenzymes OGOX1 and CELLOX and metallo-oxidoreductases (POD) can work together under the same apoplastic conditions (pH 5.5) and transduce the cell wall hydrolysis performed by microbial GHs into biochemical reactions involved both in defence and growth. This aspect may allow the plants to mount a balanced response by lowering the metabolic costs and deleterious effects deriving from an exaggerated activation of their immunity (Benedetti et al., 2015). It is also worth mentioning that H_2_O_2_ is *per se* an important transduction signal and the recent identification of the extracellular H_2_O_2_ sensor HPCA1 from *A. thaliana* reinforces its role as a cell-to-cell signal in plant immunity. Here, H_2_O_2_-mediated modification of the cysteine residues localized in HPCA1 ectodomain leads to stomatal closure, a well know defence response against pathogenic bacteria (Wu et al., 2020).

**Fig. 4.**
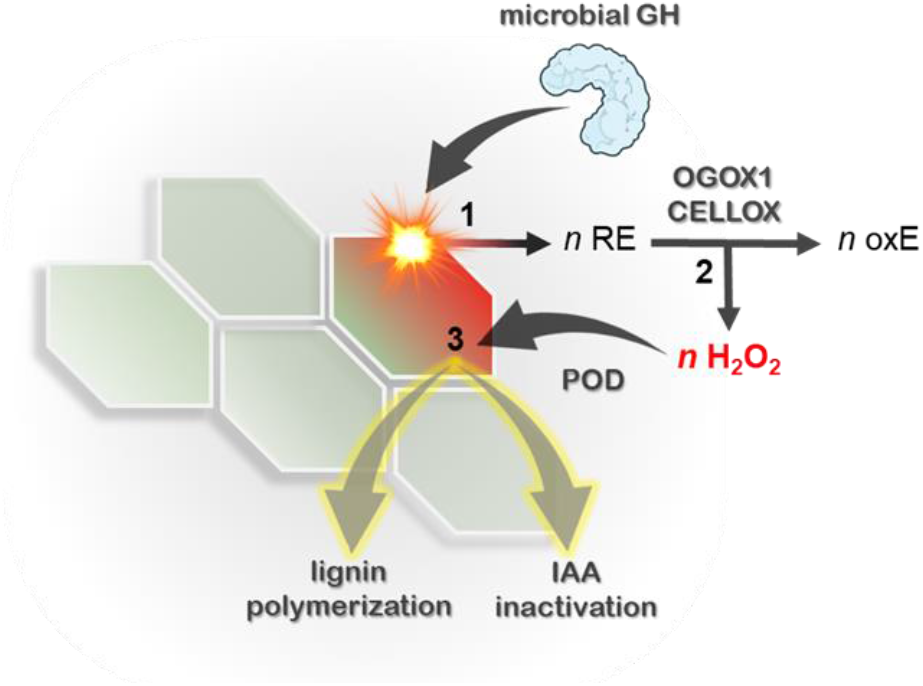
Proposed model of OGOX1/CELLOX as transducers between microbial GHs and plant PODs. The combined action of a microbial GH, a specific berberine bridge enzyme-like oligosaccharide oxidase (OGOX1, CELLOX) and a plant POD succeeded in converting the hydrolysis of a cell wall polysaccharide to lignin polymerization and auxin inactivation over degradation time. Black numbers (1-3) indicate the sequential order of the enzymatic reactions. [CELLOX: CD-oxidase from *A. thaliana*, GH: Glycoside hydrolase, IAA: indole-3-acetic acid, OGOX1: OG-oxidase 1 from *A. thaliana*, oxE: oxidized end, POD: Peroxidase, RE: reducing end].

Interestingly, oligosaccharide oxidases are also produced by phytopathogens and saprotrophs. In this case, H_2_O_2_ produced from their activity may be used by microbial lytic polysaccharide monooxygenases (LPMOs) to degrade cellulose, xylan and pectin (Villares et al., 2017; Couturier et al., 2018; Sabbadin et al., 2021) since the copper-containing active site of LPMOs can be reactivated through a H_2_O_2_-mediated reduction (Müller et al., 2018).

In conclusion, our study provides a novel perspective on how the cell wall hydrolysis can be perceived and managed by plants to balance growth and defence (Fig. 4). The high number of PODs in plants and the possible occurrence of many other BBE-l oligosaccharide oxidases in addition to OGOX1 and CELLOX poses major challenges in elucidating their role not only in plant-microbe interactions but also in plant development, morphogenesis and growth.

## ABBREVIATIONS

ABTS: 2,2’-azino-bis-(3-ethylbenzothiazoline-6-sulfonic acid)
AnEG: endoglucanase from *Aspergillus niger* (poly-α-1,4-galacturonide glycanohydrolase, EC number: 3.2.1.15)
APOD: anionic peroxidase from ripe tomato fruit (phenolic donor: hydrogen-peroxide oxidoreductase, EC number: 1.11.1.7)
BBE: Berberine Bridge Enzyme [(S)-reticuline:oxygen oxidoreductase, EC number: 1.21.3.3]
BBE-l: Berberine Bridge Enzyme-like
CAT: catalase from bovine liver (hydrogen-peroxide:hydrogen-peroxide oxidoreductase, EC number: 1.11.1.6)
CD: cellodextrin
CELLOX: cellodextrin-oxidase, CD-oxidase (cellodextrin:oxygen oxidoreductase, EC number: 1.21.3.3)
DAMP: damage-associated molecular pattern
GH: glycoside hydrolase
HRP: horseradish peroxidase VI-A type (phenolic donor: hydrogen-peroxide oxidoreductase, EC number: 1.11.1.7)
IAA: indole-3-acetic acid, auxin
OG: oligogalacturonide
OGOX: oligogalacturonide-oxidase, OG-oxidase (oligogalacturonide:oxygen oxidoreductase, EC number: 1.21.3.3)
FpPG: endopolygalacturonase from *Fusarium phyllophilum* (poly-α-1,4-galacturonide glycanohydrolase, EC number: 3.2.1.15)
POD: peroxidase (phenolic donor: hydrogen-peroxide oxidoreductase, EC number: 1.11.1.7)

## AUTHOR CONTRIBUTIONS

M.B. and B.M. conceived the project. M.B. designed the experiments, A.S. performed the experiments and analyzed the data jointly with F.A., F.C., G.D.L., M.B. and B.M; V.S. and M.G. contributed to perform the experiments. A.S., M.G. and M.B. wrote the manuscript draft whereas F.A., F.C., G.D.L., M.B. and B.M. edited the final version of the manuscript. B.M. and M.B. supervised the research. All authors have approved the final manuscript.

## ACKNOWLEDGEMENTS

The authors gratefully acknowledge Prof. Giuseppina Pitari (Dept. of Life, Health and Environmental Sciences, University of L’Aquila) for inspiring discussions on flavoenzymes.

## SUPPLEMENTARY INFORMATION

**Methods S1**. Experimental material and methods.

**Data S1**. Gene sequence used for the heterologous expression of Flag-His-SUMOstar-CELLOX (FHS-CELLOX) in *P. pastoris*.

**Table S1**. ATG code of 73 different class III PODs from *A. thaliana*.

**Fig. S1**. Heterologous expression of FHS-CELLOX in *P. pastoris*.

**Fig. S2**. Purification of the enzymes heterologously expressed in *P. pastoris*.

**Fig. S3**. H_2_O_2_-conversion efficiency of OGOX1 and FHS-CELLOX.

**Fig. S4**. Determination of ABTS-oxidizing activity of APOD from ripe tomato fruit.

**Fig. S5**. Heatmap of gene expression levels of OGOX1, CELLOX and different class III PODs from *A. thaliana*.

## Methods S1

### Design of the constructs expressing OGOX1, CELLOX and FpPG

The constructs expressing Arabidopsis OGOX1 (pPICZαB/H-OGOX1) and the polygalacturonase from *Fusarium phyllophilum* (FpPG, pGAPZαA/FpPG) were the same used in (Benedetti et al., 2011; Scortica et al., 2021), respectively. The gene encoding the mature CELLOX from *A. thaliana* (AT4G20860) was fused downstream of the SUMOstar sequence developed by LifeSensor Inc. (https://lifesensors.com/) that also included the sequences encoding the FLAG-(DYKDDDDK) and 6xHistags(HHHHHH)(https://lifesensors.com/wpcontent/uploads/2019/09/2160_2161_Pichia_SUMOstar_Manual-1.pdf). The sequence of the chimeric gene, here referred to as *FHS-CELLOX*, was codon-optimized with the codon usage of *P. pastoris* by using the online tool OPTIMIZER (http://genomes.urv.es/OPTIMIZER/) (Puigbò et al., 2007) and synthesized by Genescript (https://www.genscript.com/) by adding the restriction sites PstI and XbaI at the 5^I^ and 3^I^ ends, respectively, of the gene. The gene *FHS-CELLOX* was then cloned in pPICZαB expression vector (Invitrogen, San Diego, USA) in frame with the sequence encoding the yeast α factor for the secretion of recombinant proteins in the medium.

### Heterologous expression of OGOX1, FHS-CELLOX and FpPG in *Pichia pastoris*

OGOX1 and FpPG were heterologously expressed in *P. pastoris* by following the same procedures described in (Benedetti et al., 2011; Scortica et al., 2021), respectively. Transformation and selection of *Pichia* transformants expressing FHS-CELLOX were performed by following the same procedures described in (Scortica et al., 2021) with some modifications. In particular, to further improve the detection of FHS-CELLOX, the culture filtrates from different *Pichia* transformants were pretreated with PNGase F (New England Biolabs, Ipswich, USA) and then analyzed by immuno-decoration analysis by using a monoclonal anti-HIS antibody (AbHis, Bio-rad, Hercules, USA). The immobilized metal affinity chromatography (IMAC) was used to bind FHS-CELLOX whereas the elution was performed by using a linear gradient of imidazole. The eluted protein was dialyzed in 50 mM Tris-HCl pH 7.5 and 100 mM (NH_4_)_2_SO_4_ by using a Vivaspin 30,000 MWCO PES (Sartorius, Gottinga, Germany).

### Evaluation of H_2_O_2_-conversion efficiency of OGOX1 and FHS-CELLOX

The H_2_O_2_-conversion efficiency of OGOX1 and FHS-CELLOX was determined by the orange-xylenol assay (Benedetti et al., 2018) using 15 µM penta-galacturonic oligosaccharide (Elicityl SA, Crolles, France) or 15 µM cello-triose (Sigma-Aldrich, Saint Louis, USA), respectively, and 100 ng of each purified enzyme in a reaction volume of 0.1 mL. Values of H_2_O_2_-conversion efficiency (%) expressed the percentage ratio of µmoles of H_2_O_2_ released from µmoles of substrate reducing ends over reaction time. Determination of reducing ends of each substrate was performed according to (Lever, 1972) using different amounts of glucose as calibration curve. All the enzymatic assays were performed in 20 mM Na acetate pH 5.5 and 50 mM NaCl at 25 °C.

### Bulk extraction of anionic tomato peroxidases (APOD) and evaluation of the activity by the ABTS-POD coupled assay

The extraction of anionic tomato peroxidases (APOD) was performed according to (Andrews et al., 2002) with some modifications. In brief, 10 gr of fresh ripe tomato fruit were frozen in liquid nitrogen and homogenized in a MM500 VARIO Mixer Mill (Retsch, Basel, Switzerland) by using a 25 mL screw-top grinding jars containing one grinding ball (15 mm) for 3-5 min at 30 Hz. The homogenized tissue was resuspended in 20 mL of a buffer composed of 50 mM Na acetate pH 5.0 and 0.5 M NaCl and incubated at 4°C under gentle shaking for 1 hour. The suspension was centrifuged at 2800 x g for 20 min and the supernatant filtered using a PES Syringe filter (0.2 µm). The filtrate was dialyzed and concentrated (16X) using a Vivaspin 10,000 MWCO PES (Sartorius, Gottinga, Germany) and quantified by the Bradford reagent (Bio-rad, Hercules, USA). The sample prepared according to this procedure was referred to as APOD. The sample was tested for the capability of oxidizing ABTS in the presence of 50 µM H_2_O_2_ (ABTS-POD coupled assay) using a reaction buffer composed of 100 µM ABTS 2,2’-azino-bis-(3-ethylbenzothiazoline-6-sulfonic acid) (Sigma-Aldrich, Saint Louis, USA) and 0.14 g.L^-1^ APOD (5% v/v, 100 μL total volume). Enzyme activity was spectrophotometrically determined at 25°C by using an Infinite® M Nano200 spectrophotometer (Tecan AG, Männedorf, Switzerland). The oxidation of ABTS to the cationic radical ABTS^+•^ was measured in continuum mode for 25 min at 415 nm (ε_415nm_ = 34 mM^-1^ cm^-1^).

### Tetra-guaiacol polymerization

The oxidative polymerization of guaiacol to tetra-guaiacol was measured by following the increase in absorbance at 470 nm (ε_470nm_ = 26.6 mM^-1^ cm^-1^) (Koduri and Tien, 1995). The OGOX1/(FHS-)CELLOX-HRP assay was performed in 20 mM Na acetate pH 5.5 containing 0.5% (w/v) polygalacturonic acid (Sigma-Aldrich, Saint Louis, USA) or 0.5% (w/v) carboxy-methyl cellulose (P-CMC4M; Megazyme, Dublin, Ireland), 150 µM guaiacol [2-methoxyphenol, (Sigma-Aldrich, Saint Louis, USA)] and 0.05 g.L^-1^ horseradish peroxidase VI-A type (HRP) (Sigma-Aldrich, Saint Louis, USA) in a reaction volume of 0.2 mL. The mixture also included a GH enzyme [7 mg.L^-1^ FpPG or 0.2 mg.L^-1^ endoglucanase from *Aspergillus niger* (AnEG) (E-CELAN; Megazyme, Dublin, Ireland)] and the appropriate BBE-l oligosaccharide oxidase (3 mg.L^-1^ OGOX1 or 3 mg.L^-1^ FHS-CELLOX). To assess the involvement of H_2_O_2_ in the oxidative polymerization of guaiacol, a catalase (CAT) from bovine liver (Sigma-Aldrich, Saint Louis, USA) was added to the reaction (0.02 g.L^-1^). The activity of the OGOX1/(FHS-)CELLOX-HRP machinery was spectrophotometrically measured at 25°C by using an Infinite® M Nano200 spectrophotometer (Tecan AG, Männedorf, Switzerland) in continuum mode for 60 min.

### IAA oxidation

IAA oxidation was measured using the modified Salkowski method described in (Gang et al., 2019). The OGOX1/(FHS-)CELLOX-APOD assay was performed in 20 mM Na acetate pH 5.5 containing 0.5% (w/v) polygalacturonic acid (Sigma-Aldrich, Saint Louis, USA) or 0.5% (w/v) carboxy-methyl cellulose (P-CMC4M; Megazyme, Dublin, Ireland), 500 µM IAA [indole-3-acetic acid, auxin (Sigma-Aldrich, Saint Louis, USA)] and 0.14 g.L^-1^ APOD in a reaction volume of 0.1 mL. The mixture also included a GH enzyme (7 mg.L^-1^ FpPG or 0.2 mg.L^-1^ AnEG) and the appropriate BBE-l oligosaccharide oxidase (3 mg.L^-1^ OGOX1 or 3 mg.L^-1^ FHS-CELLOX). IAA oxidation was measured at 25°C by following the decrease in absorbance at 536 nm using an Infinite® M Nano200 spectrophotometer (Tecan AG, Männedorf, Switzerland). Each absorption value was converted in µM IAA by interpolation with the IAA-calibration curve and then converted to percentage of residual IAA (% IAA) respect to the starting concentration (i.e., 500 µM corresponds to 100% IAA).

---

**Data S1.**
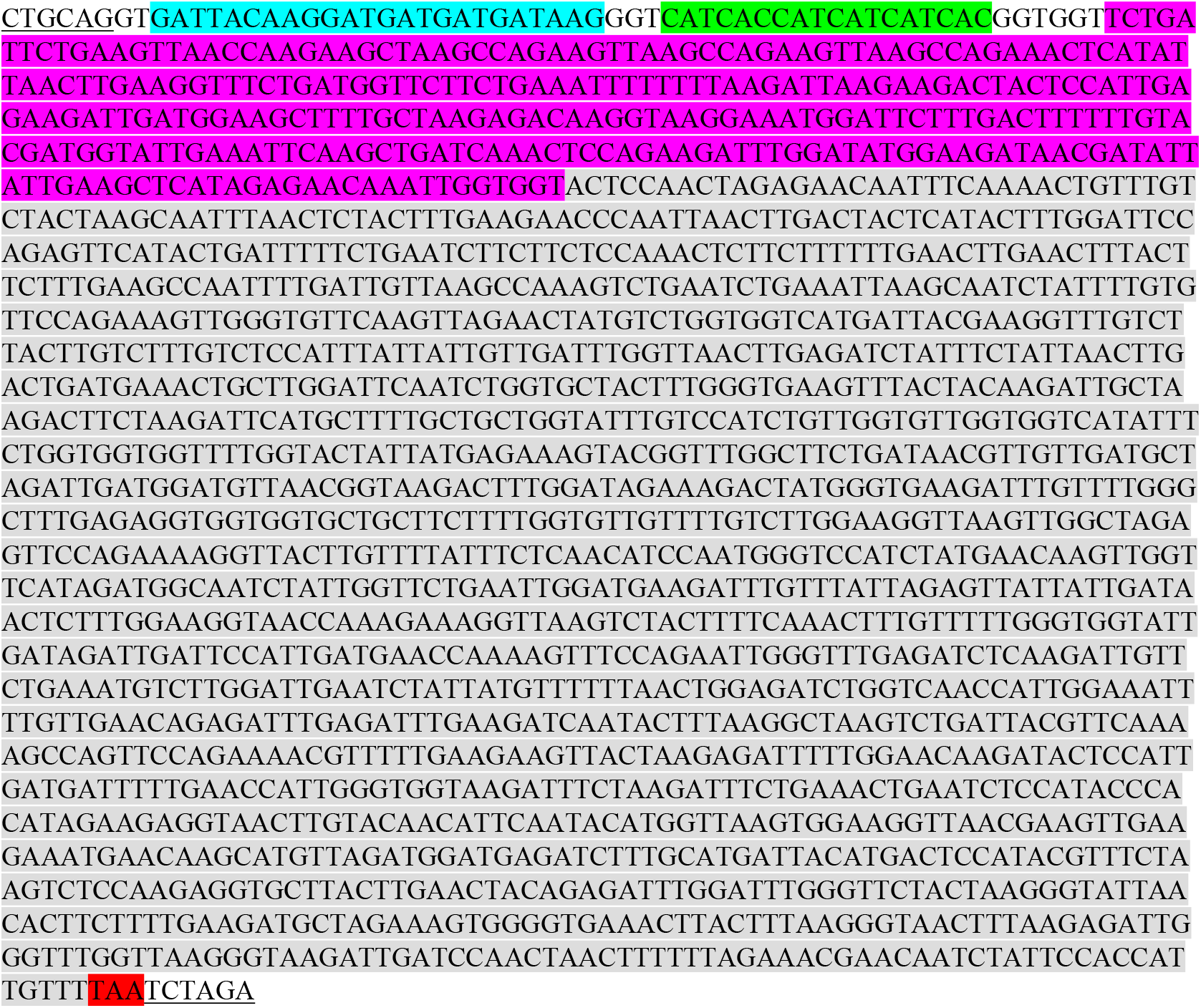
Gene sequence used for the heterologous expression of Flag-His-SUMOstar-CELLOX (FHS-CELLOX) in *P. pastoris*. Codon-optimized sequence of FHS-CELLOX used for the expression in *Pichia pastoris*. The sequence was fused downstream of the sequence encoding the α-factor signal peptide from the vector pPICZαB. Underlined sequences: restriction sites used for cloning (i.e., PstI and XbaI); turquoise sequence: Flag epitope-encoding sequence; green sequence: 6xHis tag-encoding sequence; purple sequence: codon-optimized sequence encoding the SUMOstar tag; grey sequence: codon-optimized sequence encoding the mature CELLOX; red sequence: STOP codon.

**Table S1.** ATG code of 73 different class III PODs from *A. thaliana*. In the table, the ATG codes of the Arabidopsis class III PODs (PER1-73) used in the heatmap (Fig. S5) are reported.

| ATG code | Gene name |
| --- | --- |
| AT1G05240 | *PER1* |
| AT1G05250 | *PER2* |
| AT1G05260 | *PER3* |
| AT1G14540 | *PER4* |
| AT1G14550 | *PER5* |
| AT1G24110 | *PER6* |
| AT1G30870 | *PER7* |
| AT1G34510 | *PER8* |
| AT1G44970 | *PER9* |
| AT1G49570 | *PER10* |
| AT1G68850 | *PER11* |
| AT1G71695 | *PER12* |
| AT1G77100 | *PER13* |
| AT2G18140 | *PER14* |
| AT2G18150 | *PER15* |
| AT2G18980 | *PER16* |
| AT2G22420 | *PER17* |
| AT2G24800 | *PER18* |
| AT2G34060 | *PER19* |
| AT2G35380 | *PER20* |
| AT2G37130 | *PER21* |
| AT2G38380 | *PER22* |
| AT2G38390 | *PER23* |
| AT2G39040 | *PER24* |
| AT2G41480 | *PER25* |
| AT2G43480 | *PER26* |
| AT3G01190 | *PER27* |
| AT3G03670 | *PER28* |
| AT3G17070 | *PER29* |
| AT3G21770 | *PER30* |
| AT3G28200 | *PER31* |
| AT3G32980 | *PER32* |
| AT3G49110 | *PER33* |
| AT3G49120 | *PER34* |
| AT3G49960 | *PER35* |
| AT3G50990 | *PER36* |
| AT4G08770 | *PER37* |
| AT4G08780 | *PER38* |
| AT4G11290 | *PER39* |
| AT4G16270 | *PER40* |
| AT4G17690 | *PER41* |
| AT4G21960 | *PER42* |
| AT4G25980 | *PER43* |
| AT4G26010 | *PER44* |
| AT4G30170 | *PER45* |
| AT4G31760 | *PER46* |
| AT4G33420 | *PER47* |
| AT4G33870 | *PER48* |
| AT4G36430 | *PER49* |
| AT4G37520 | *PER50* |
| AT5G05340 | *PER51* |
| AT4G37530 | *PER52* |
| AT5G06720 | *PER53* |
| AT5G06730 | *PER54* |
| AT5G14130 | *PER55* |
| AT5G15180 | *PER56* |
| AT5G17820 | *PER57* |
| AT5G19880 | *PER58* |
| AT5G19890 | *PER59* |
| AT5G22410 | *PER60* |
| AT5G24070 | *PER61* |
| AT5G39580 | *PER62* |
| AT5G40150 | *PER63* |
| AT5G42180 | *PER64* |
| AT5G47000 | *PER65* |
| AT5G51890 | *PER66* |
| AT5G58390 | *PER67* |
| AT5G58400 | *PER68* |
| AT5G64100 | *PER69* |
| AT5G64110 | *PER70* |
| AT5G64120 | *PER71* |
| AT5G66390 | *PER72* |
| AT5G67400 | *PER73* |

**Fig. S1.**
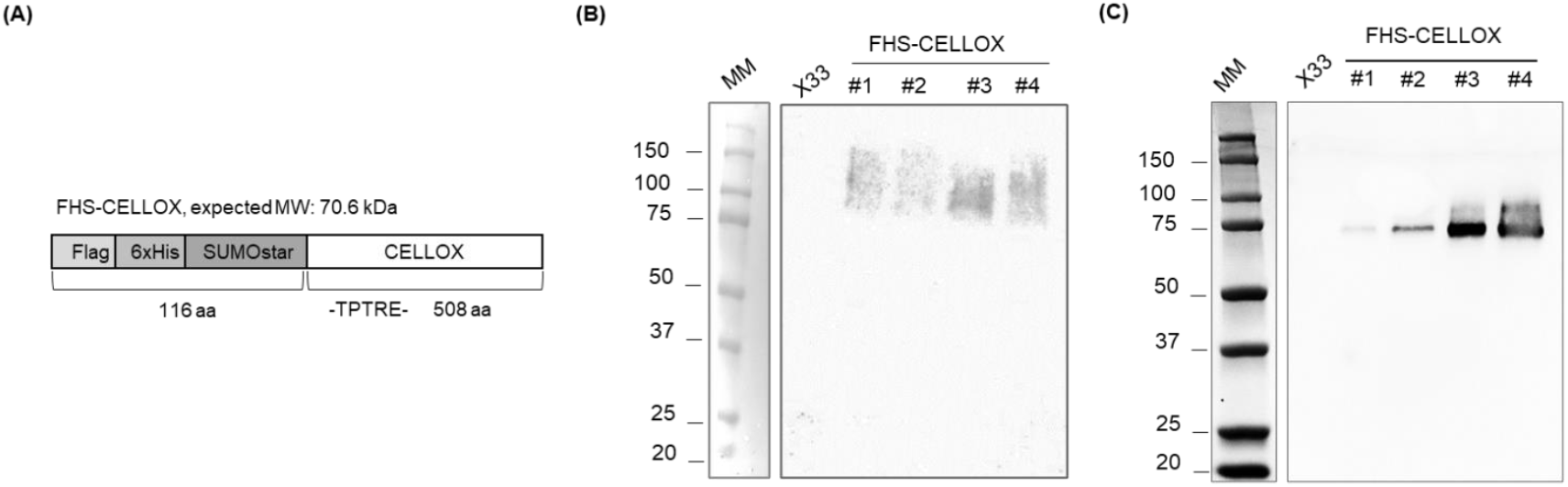
Heterologous expression of FHS-CELLOX in *P. pastoris*. (A) Schematic representation of Flag-His-SUMOstar-tagged CELLOX, referred to as FHS-CELLOX. In the scheme, the starting amino acid sequence of mature CELLOX (-TPTRE-) was fused downstream of Flag-His-SUMOstar-tag. (B) Immuno-decoration analysis of the raw cultures filtrates from four different (#1-4) *P. pastoris* transformants expressing FHS-CELLOX. (C) Immuno-decoration analysis of the same culture filtrates shown in (B) upon deglycosylation with PNGase F. The (B) raw and (C) PNGase F-treated culture filtrates of wild type *P. pastoris* (X33) were used as negative controls. Molecular weight marker (MM) is also reported.

**Fig. S2.**
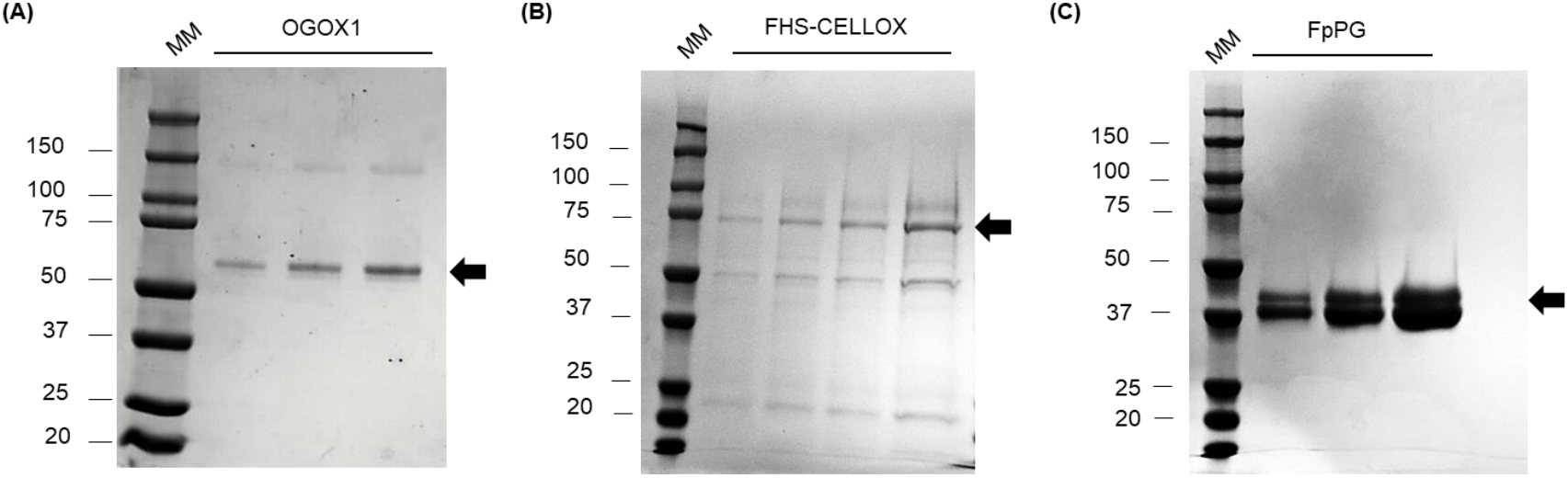
Purification of the enzymes heterologously expressed in *P. pastoris*. SDS-PAGE/Coomassie blue staining analysis of different amounts of purified (A) OGOX1, (B) FHS-CELLOX upon deglycosylation with PNGase F and (C) FpPG. (A-C) Black arrows point to the bands corresponding to (A) OGOX1, (B) FHS-CELLOX and (C) FpPG. Molecular weight marker (MM) is also reported. [FHS-CELLOX: Flag-His-SUMOstar-tagged CD-oxidase from *A. thaliana*, FpPG: endopolygalacturonase from *F. phyllophilum*, OGOX1: His-tagged OG-oxidase 1 from *A. thaliana*].

**Fig. S3.**
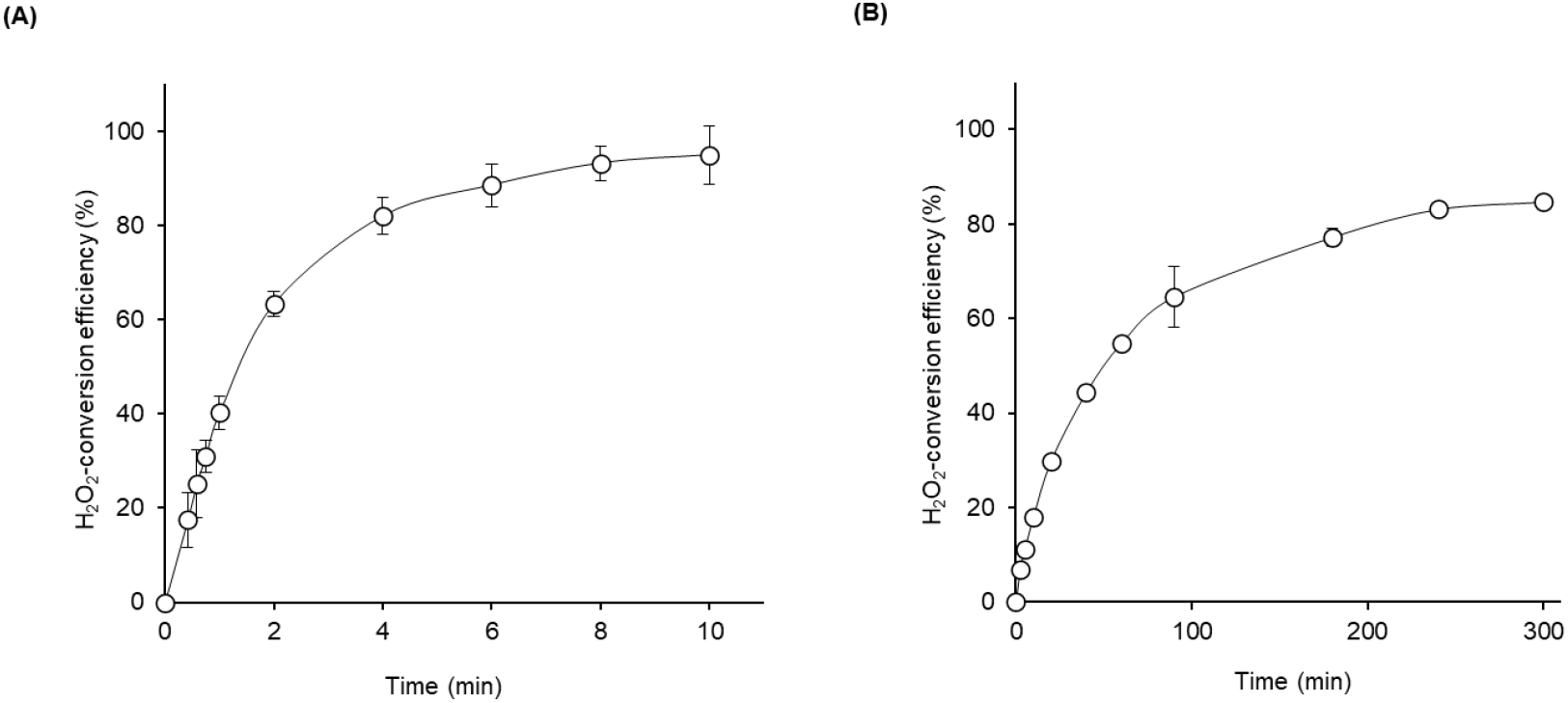
H_2_O_2_-conversion efficiency of OGOX1 and FHS-CELLOX. The H_2_O_2_-conversion efficiency of (A) OGOX1 and (B) FHS-CELLOX was evaluated by measuring the production of H_2_O_2_ in the presence of 15 µM penta-galacturonic oligosaccharide or cello-triose, respectively, by using the orange xylenol assay. Values are mean ± s.d. (n= 3). [FHS-CELLOX: Flag-His-SUMOstar-tagged CD-oxidase from *A. thaliana*, OGOX1: His-tagged OG-oxidase 1 from *A. thaliana*].

**Fig. S4.**
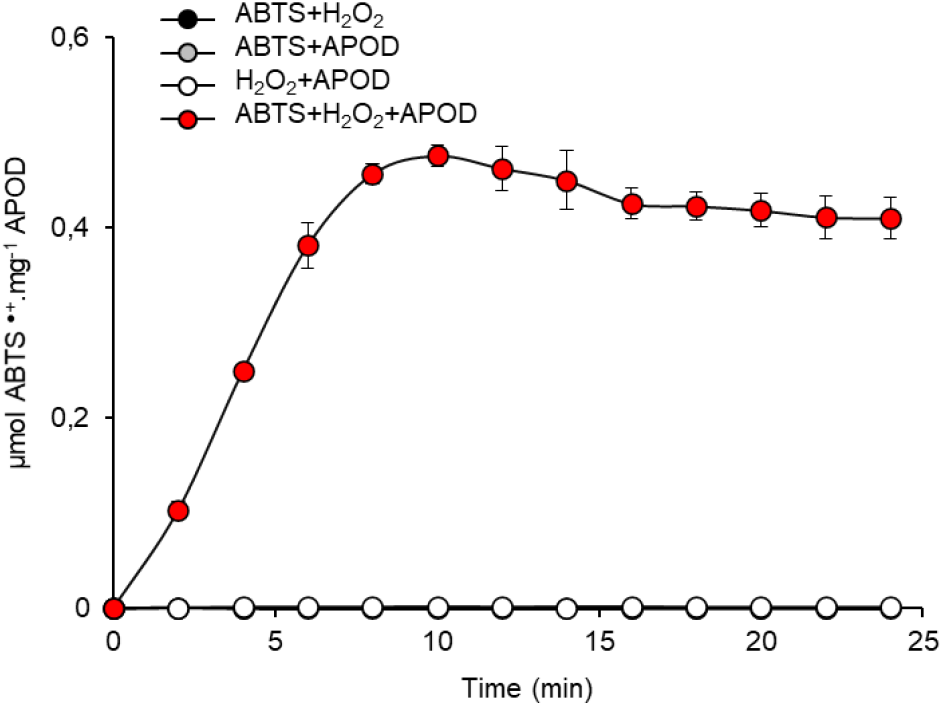
Determination of ABTS-oxidizing activity of APOD from ripe tomato fruit. ABTS-oxidizing activity of an anionic peroxidase preparation from ripe tomato fruit extract (APOD) in the presence of 50 µM H_2_O_2_ as determined by ABTS-APOD coupled assay. Values are mean ± s.d. (n= 3). The kinetics relative to the samples (ABTS+H_2_O_2_) and (ABTS+APOD) superpose with that of the sample (H_2_O_2_+APOD). [ABTS: 2,2’-azino-bis-(3-ethylbenzothiazoline-6-sulfonic acid), APOD: anionic peroxidase preparation from ripe tomato fruit].

**Fig. S5.**
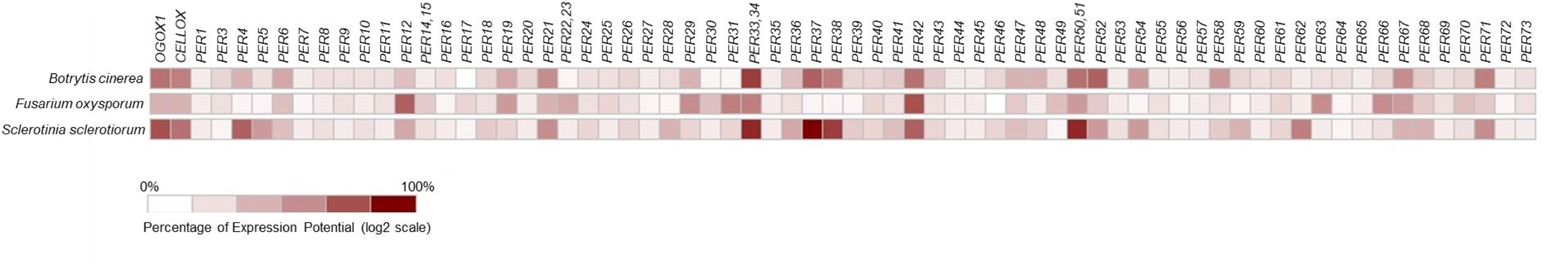
Heatmap of gene expression levels of OGOX1, CELLOX and different class III PODs from *A. thaliana*. The columns of the heatmap represent genes (OGOX1-, CELLOX- and different class III POD-encoding genes) whereas the rows represent the experimental conditions (three different fungal infections on *A. thaliana* Col-0, n= 1, 48h post-inoculation). Each cell is colorized in different red intensities based on the level of expression of that gene in that sample. *PER2* is not available. Further details on plant PODs (PER1-73) used in the analysis are reported in Table S1. Heatmap has been created by https://genevestigator.com/. [CELLOX: CD-oxidase, AT4G20860; OGOX1: OG-oxidase 1, AT4G20830; POD: Peroxidase].

